# GPR182 promotes atherosclerosis via facilitating arterial lipid deposition

**DOI:** 10.64898/2026.09.15.748866

**Authors:** Taichi Terai, Shunsuke Doi, Zhiwei Sun, Yi Sun, Dustin P Fykstra, Yujie Guo, Yuya Hirasawa, Junyi Hu, Masayuki Sho, Richard D Schulick, Yuwen Zhu

## Abstract

Elevated circulating low-density lipoprotein cholesterol (LDL-C) is a primary risk factor for atherosclerosis; however, the molecular mechanisms underlying lipid plaque formation within the artery wall remain poorly understood. Here, we identify GPR182, a recently characterized lipoprotein receptor of the atypical chemokine receptor family, as a key mediator of lipid deposition in the aortic endothelium during atherosclerosis. Genetic depletion of GPR182 protects against atherosclerosis across multiple mouse models without altering circulating cholesterol levels or immune cell recruitment to plaques. GPR182 is expressed by aortic endothelial cells (ECs) and is further upregulated during disease progression. GPR182 mediates LDL uptake by aortic ECs both in vitro and in vivo. Blockade of GPR182 with a monoclonal antibody reduces lipid uptake in the aorta and attenuates disease progression under hypercholesterolemic conditions. Collectively, these findings identify endothelial GPR182 as a critical regulator of aortic lipid deposition and atherosclerotic plaque formation and support GPR182 inhibition as a promising therapy for atherosclerotic cardiovascular disease.

## Introduction

Atherosclerotic cardiovascular disease remains a leading cause of mortality worldwide. Elevated circulating LDL cholesterol (LDL-C) is closely associated with the risk of cardiovascular diseases^1^, and therapies that lower plasma LDL-C effectively reduce cardiovascular risks^2–4^. However, significant residual cardiovascular risk persists even in patients achieving very low LDL-C levels^5^, highlighting the need for therapeutic strategies that complement systemic lipid lowering. Entry of circulating LDL into the arterial wall is a critical early event in atherosclerosis, and emerging evidence indicates that LDL transport through the arterial endothelium is an active and highly regulated process involving receptor-mediated uptake and transcytosis. Several cell-surface receptors expressed by arterial endothelial cells (AECs), including scavenger receptor class B type I (SR-BI) and activin receptor-like kinase 1 (ALK1), have been implicated in LDL binding, internalization, and/or transendothelial transport^6–8^. Genetic deletion or antibody-mediated blockade of these receptors reduces arterial lipid accumulation and atherosclerosis in experimental mouse models without necessarily altering circulating lipoprotein levels^9^. Thus, characterizing the molecular machinery that controls endothelial lipoprotein transport may reveal therapeutic targets that directly limit lipid deposition within the arterial wall and complement existing systemic cholesterol-lowering therapies.

GPR182, also known as ACKR5^10^, is a recently characterized member of the atypical chemokine receptor (ACKR) family^11^ ^12^. GPR182 is predominantly expressed by ECs, including lymphatic and sinusoidal endothelial cells as well as vascular endothelial cells in multiple tissues^11,13,14^. Compared with other ACKRs, GPR182 interacts with a broader spectrum of chemokines^12,14,15^ ^16^. We recently identified GPR182 as a lipoprotein receptor that regulates dietary lipid absorption^17^, raising the possibility that GPR182 may also participate in endogenous lipoprotein transport. Here, we show that GPR182 is constitutively expressed by aortic ECs and is further upregulated during atherosclerosis. Our investigations further demonstrate that GPR182 mediates LDL uptake and lipid deposition within the aortic wall, and that GPR182 ablation inhibits atherosclerotic plaque development.

## Results

### GPR182 deficiency attenuates atherosclerosis in *Apoe*−/− mice

To evaluate the role of GPR182 in atherosclerosis, we generated *Gpr182*−/−*Apoe*−/− mice to assess atherosclerotic lesion development. At 10-months-old, male and female *Apoe*−/− mice on a regular chow diet developed extensive aortic atherosclerosis, whereas age-matched *Gpr182*−/−*Apoe*−/− mice exhibited markedly reduced plaque burden (**Figure S1A**). En face Oil Red O staining confirmed reduced aortic lesion area in *Gpr182*−/−*Apoe*−/− mice (**Figure S1B**). Consistent with these findings, histological analysis of the aortic root revealed significantly smaller lesions in *Gpr182*−/−*Apoe*−/− mice (**Figure S1C**); Atherosclerotic plaques from *Gpr182*−/−*Apoe*−/− mice had less lipid content than those from *Apoe*−/− controls (**Figure S1D**).

We next assessed atherosclerosis progression in young adult mice challenged with a high-cholesterol diet (HCD) for 10 weeks. Body weight gain was comparable between *Apoe*−/− and *Gpr182*−/−*Apoe*−/− mice in both sexes (**Figure 1A**). In situ visualization of the aorta in *Gpr182*−/− *Apoe*−/− mice revealed a substantial reduction in plaque burden. Atherosclerotic plaques occupied >20% of the aortic arch in *Apoe*−/− mice compared with 10% in *Gpr182*−/−*Apoe*−/− mice (**Figure 1B**). En face Oil Red O staining demonstrated over 50% reduction in aortic lesion area in *Gpr182*−/−*Apoe*−/− mice (**Figure 1C**), and cross-sectional analysis of the aortic root further confirmed reduced lesion area (**Figure 1D**). Compared with *Apoe*−/− controls, lesions from *Gpr182*−/−*Apoe*−/− mice exhibited reduced lipid accumulation (**Figure 1E**), accompanied b increased collagen content (**Figure 1F**). Consistently, ApoB staining was markedtly decreased in aortic lesions from *Gpr182*−/−*Apoe*−/− mice (**Figure 1G**). Together, these findings demonstrate that GPR182 deficiency reduces atherosclerotic lesion burden and is associated with a plaque phenotype characterized by less lipid and more collagen, consistent with a more stable lesion composition.

**Figure 1.**
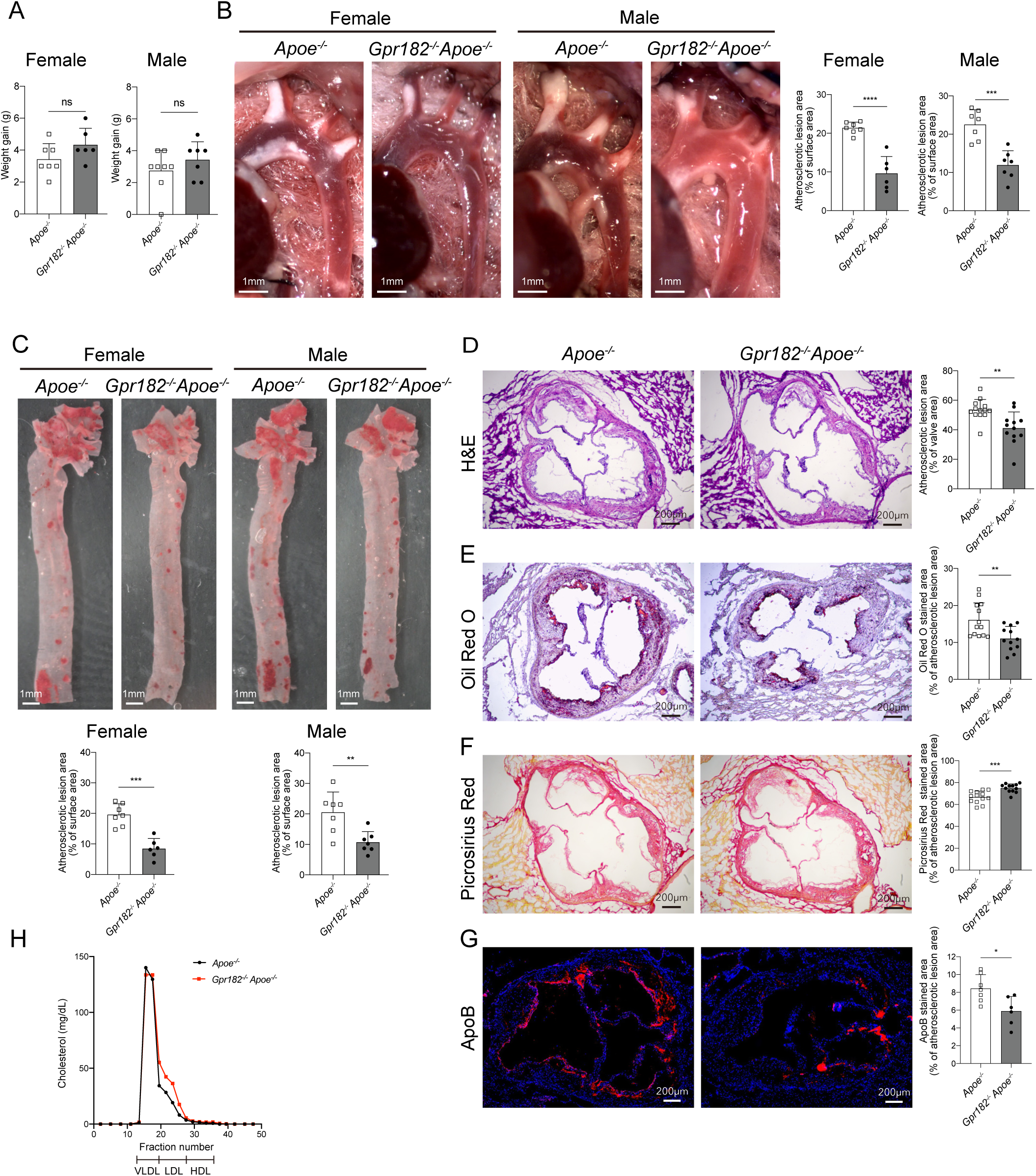
GPR182 ablation attenuates atherosclerosis progression in *Apoe*−/− mice under HCD. Young adult male and female *Apoe*−/− and *Gpr182*−/−*Apoe*−/− mice were fed an HCD for 10 weeks. (A) Body weight gain after feeding was recorded. (B) Representative in situ aortic arch image of atherosclerotic plaque, with quantification of lesion area. (C) Representative Oil Red O stained en face images of aortas, with quantification of lesion area. (D) H&E-stained sections of aortic root, with lesion area quantification. (E) Oil Red O staining of aortic root, with lipid content quantification. (F) Picrosirius red staining of aortic root, with collagen content quantification. (G) ApoB staining in aortic root lesions, with quantification of ApoB-positive area. (H) FPLC fractionation was performed to assess serum cholesterol distribution across lipoprotein fractions.

Because GPR182 functions as a lipoprotein receptor^17^, we next determined whether the reduction in atherosclerosis was associated with changes in circulating lipid levels. On a regular chow diet, serum triacylglycerol (TAG) and total cholesterol levels were comparable between *Apoe*−/− and *Gpr182*−/−*Apoe*−/− mice (**Table S1**). After 10 weeks of HCD feeding, serum cholesterol increased to a similar extent in both groups (**Table S1**). Fast protein liquid chromatography (FPLC) analysis of serum lipoproteins further revealed comparable cholesterol profiles across VLDL, LDL, and HDL fractions between *Apoe*−/− and *Gpr182*−/−*Apoe*−/− mice (**Figure 1H**). Serum triglyceride and total cholesterol levels were also similar between the two genotypes of aged mice (**Table S1**). Thus, GPR182 deficiency reduces atherosclerosis without impacting circulating cholesterol levels.

### *Gpr182*−/− mice are protected from AAV8-PCSK9-induced atherosclerosis

To determine whether the atheroprotective effect of GPR182 deletion extends to another mouse model of atherosclerosis, we employed the AAV8-PCSK9 model. In this model, a single injection of AAV8 encoding gain-of-function PCSK9 drives hepatic LDL receptor degradation, inducing hypercholesterolemia and atherosclerosis on an intact C57BL/6J background, without requiring apoE deficiency ^18^. Male *Gpr182*−/− mice and age-matched wild-type (WT) controls were intravenously administered AAV8-PCSK9 and subsequently fed an HCD for 12 weeks. *Gpr182*−/− mice gained slightly less body weight than WT controls (**Figure 2A**), consistent with impaired dietary fat absorption in *Gpr182*−/− mice^17^. Serum TAG levels were lower in *Gpr182*−/− mice (**Figure 2B**); In contrast, serum cholesterol levels were comparable between the two groups of mice (**Figure 2B**). Assessment of aortic atherosclerosis demonstrated markedly reduced lesion burden in *Gpr182*−/− mice. Atherosclerotic lesion area at the aortic valve region was reduced by about 50% in *Gpr182*−/− mice compared with WT controls (**Figure 2C**, **2D**). Consistently, cross-sectional analysis of the aortic root demonstrated substantially smaller atherosclerotic lesions in *Gpr182*−/− mice (**Figure 2E**). Lesions from *Gpr182*−/− mice had significantly reduced lipid accumulation (**Figure 2F**, **2G**). Thus, the findings in the AAV8-PCSK9 model further confirms that GPR182 deficiency attenuates atherosclerosis despite comparable circulating total cholesterol levels.

**Figure 2.**
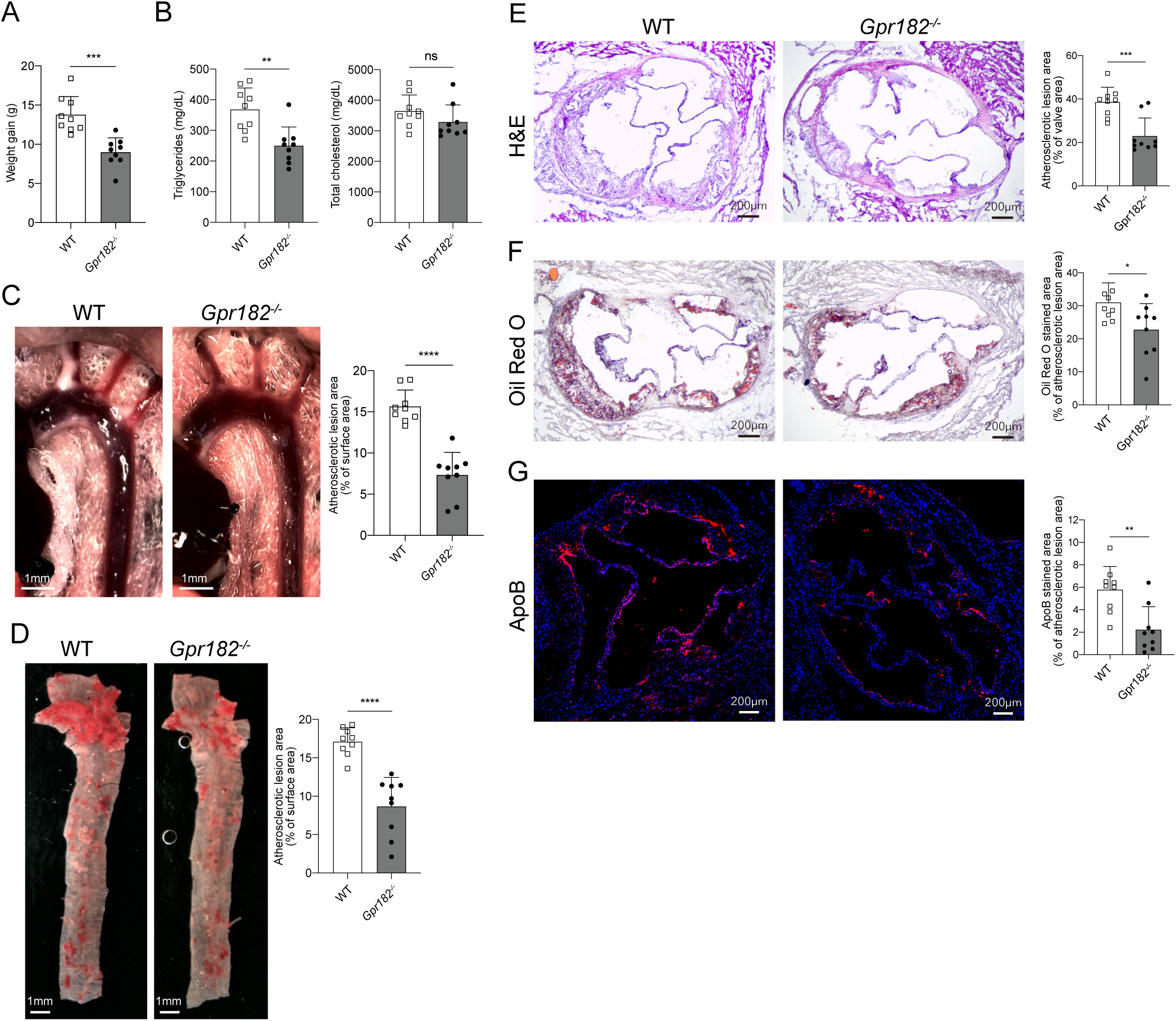
*Gpr182*−/− mice are resistant to AAV-mPCSK9 induced atherosclerosis. Male WT and *Gpr182*−/− mice were intravenously injected with AAV8-mPCSK9, followed with 12 weeks of HCD. (A) Body weight gain during HCD feeding was recorded. (B) Serum TAG and total cholesterol were quantified. (C) Representative in situ aortic arch image of atherosclerotic plaque, with quantification of lesion area. (D) Representative Oil Red O stained en face images of aortas, with quantification of lesion area. (E) H&E-stained sections of aortic root, with lesion area quantification. (F) Oil Red O staining of aortic root, with lipid content quantification. (G) ApoB staining in aortic root lesions, with quantification of ApoB-positive area.

### GPR182 is enriched in aortic endothelial cells

Because GPR182 deficiency attenuated atherosclerosis in multiple mouse models without altering circulating cholesterol levels, we investigated whether GPR182 is expressed within the aortic vasculature. Analysis of the Tabula Muris single-cell RNA-sequencing dataset revealed enrichment of Gpr182 expression in endocardial and vascular EC populations of the heart and aorta (**Figure S2A, S2B**). Immunofluorescence staining of mouse aortic tissues using a GPR182 mAb^17^ confirmed prominent GPR182 expression in the endothelium, as evidenced by its colocalization with the endothelial marker CD31 (**Figure 3A**). Moreover, in *Apoe*−/− mice fed an HCD, the proportion of GPR182-expressing AECs was significantly increased, as determined by flow cytometry and immunofluorescence analyses (**Figure 3B**, **S2C**).

**Figure 3.**
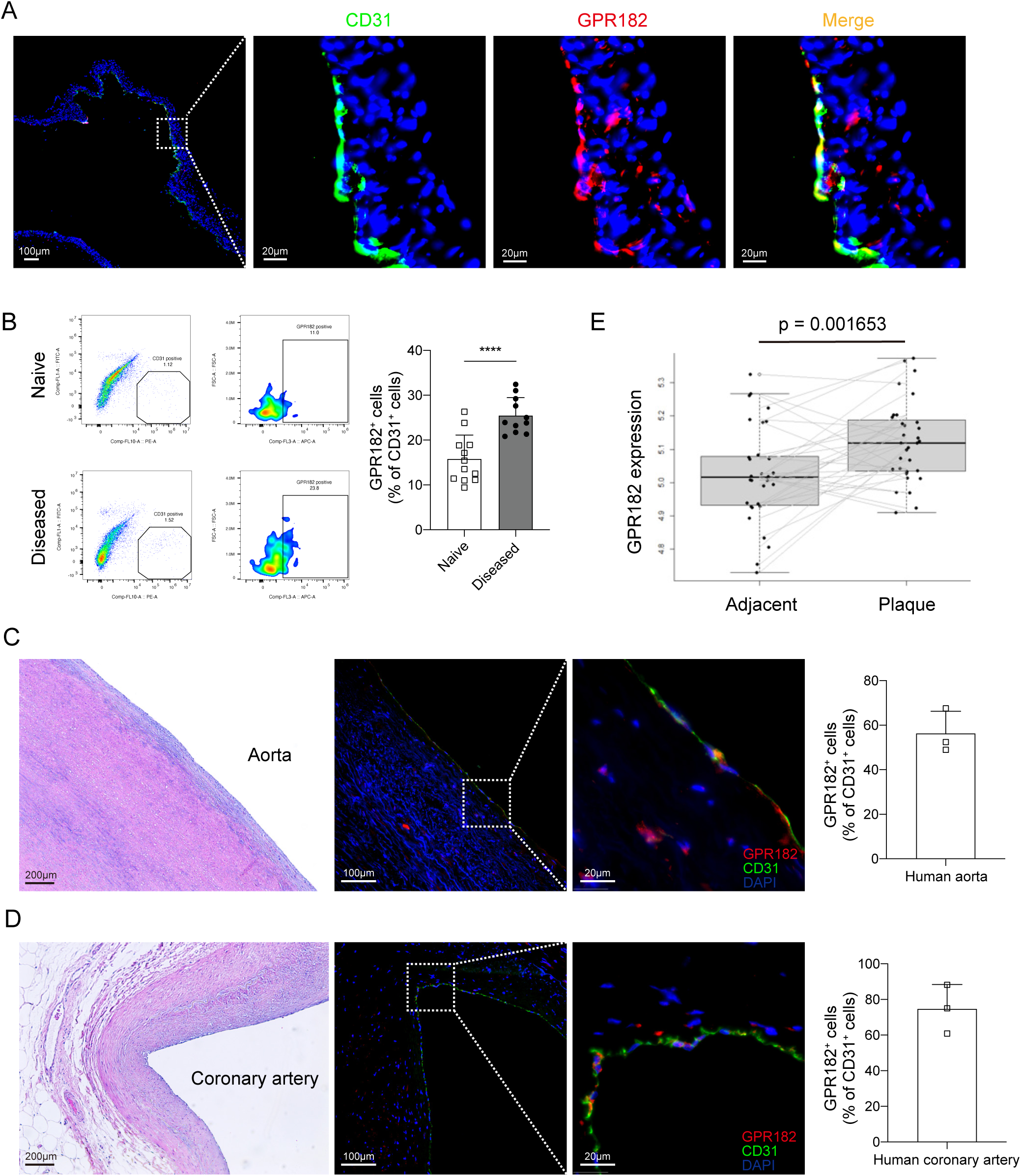
GPR182 is expressed in arterial endothelial cells and increases during atherosclerosis. (A) Immunofluorescent staining of GPR182 and CD31 in aortic tissues from naïve mice. (B) Single cell suspensions of aortic tissues from naïve and diseased mice (*Apoe*−/− mice after 10 weeks of HCD) were examined for GPR182 expression in ECs (CD31+) by flow cytometry. Immunofluorescent staining of GPR182 and CD31 in human aorta (C) and coronary artery (D). (E) GPR182 expression in atherosclerotic plaques and paired adjacent normal tissues from a cohort of 34 patients (GSE43292).

We next examined whether GPR182 is similarly expressed in human cardiovascular vessels. Nonlesional vascular tissues obtained from three donors undergoing aortic aneurysm repair or cardiac explantation (**Figure 3C**) exhibited a strong luminal GPR182 signal that colocalized with CD31 (**Figure 3D**). To determine whether vascular GPR182 expression changes during atherosclerosis, we analyzed a publicly available transcriptomic dataset of paired atherosclerotic and adjacent nonatherosclerotic aortic tissues^9^. GPR182 expression was significantly higher in atherosclerotic lesions than in paired adjacent nonatherosclerotic regions (**Figure 3E**). Together, these findings identify GPR182 as an endothelial-enriched receptor in the mouse and human cardiovascular vasculature and demonstrate increased vascular GPR182 expression in association with atherosclerosis.

### GPR182 deficiency does not alter immune cell accumulation in atherosclerotic lesions

Recruitment and accumulation of inflammatory immune cells, particularly macrophages and CD4+ T cells, are key features of atherosclerosis progression^19^ ^20,21^. As a broadly scavenging ACKR capable of engaging with multiple chemokines^12,14^, vascular GPR182 could potentially modulate local chemokine availability and thereby influence immune cell recruitment during atherogenesis. To assess this possibility, we quantified chemokines in aortic tissue extracts from *Apoe*−/− and *Gpr182*−/−*Apoe*−/− mice after 10 weeks of HCD feeding. Contrary to this expectation, GPR182 deficiency did not increase chemokine abundance in the diseased aorta. Instead, several chemokines, including CCL3 and CXCL1, were significantly reduced in aortic tissues from *Gpr182*−/−*Apoe*−/− mice compared with *Apoe*−/− controls (**Figure S3A**). Thus, the reduction of chemokines reflects the diminished overall lesion and lipid burden in *Gpr182*−/−*Apoe*−/− aortas rather than a direct effect of GPR182 on chemokine scavenging.

We next assessed immune cell accumulation within atherosclerotic lesions. Immunofluorescence analysis of the aortic valve region revealed comparable infiltration of CD45+ leukocytes and F4/80+ macrophages in *Apoe*−/− and *Gpr182*−/−*Apoe*−/− mice (**Figure S3B**). Consistent with these findings, flow cytometric analysis of single-cell suspensions prepared from whole aortas showed no significant differences in the numbers of infiltrating T cells or macrophages between the two groups (**Figure S3C**). Thus, GPR182 deficiency does not substantially affect T-cell or macrophage accumulation in the atherosclerotic aorta, suggesting that altered immune cell recruitment is unlikely to account for the attenuated atherosclerosis observed in *Gpr182*−/−*Apoe*−/− mice.

### GPR182 mediates LDL deposition in the aorta

Receptor-mediated LDL entry into the arterial wall has recently been recognized as critical to the initiation and progression of atherosclerosis ^9^. Our recent finding has established GPR182 as a lipoprotein receptor that mediates dietary lipid absorption^17^. Thus, GPR182 in AECs could contribute to the development of atherosclerosis through the uptake of circulating LDL. We injected Dil-LDL intravenously into WT and *Gpr182*−/− mice to assess whether GPR182 in the aorta mediates LDL uptake. In WT mice, injected LDL was primarily present within the CD31+ endothelium of the aorta (**Figure 4A**) and colocalized with GPR182 (**Figure 4B**). Significant LDL uptake was detected in the aorta walls of WT mice, whereas signal was barely detectable in *Gpr182*−/− aortic tissues (**Figure 4C**). A similar finding was observed in *Gpr182*−/−*Apoe*−/− mice when Dil-LDL was administered (**Figure S4A**). In contrast, *Ldlr*−/− mice exhibited normal LDL retention in the aorta comparable to WT mice, confirming the earlier report that LDLR is not involved in LDL deposition by aortic ECs *in vivo* (**Figure 4D**)^8^. Importantly, the impaired LDL uptake in *Gpr182*−/− mice was not due to altered endothelial permeability, because *Apoe*−/− mice and *Gpr182*−/−*Apoe*−/− mice under HCD exhibited similar Evans blue uptake in the aorta upon intravenous administration (**Figure S4B**).

**Figure 4.**
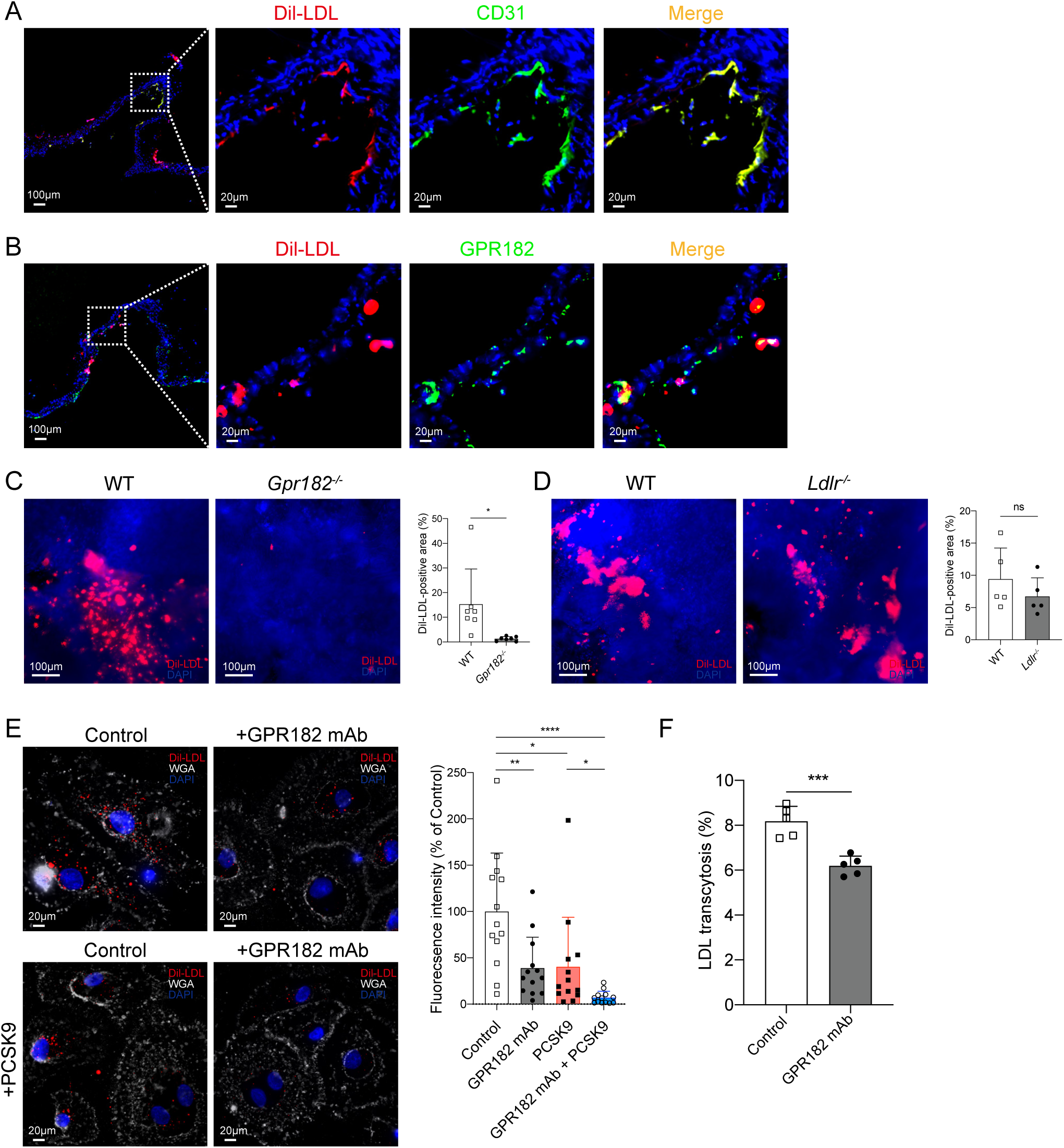
GPR182 mediates endothelial LDL uptake in the aorta. (A, B) WT C57BL/6 mice were intravenously injected with Dil-LDL. 30 minutes later, aortic tissues were collected to determine fluorescent signal. CD31 (A) or GPR182 (B) was co-stained. (C) Dil-LDL signal was detected in aorta from WT and *Gpr182*−/− mice following intravenous Dil-LDL injection, with quantification of Dil-LDL signal. (D) Dil-LDL signal was detected in aorta from WT and *Ldlr*−/− mice following intravenous Dil-LDL injection, with quantification of Dil-LDL signal. (E) HAECs with or without PCSK9 treatment were assessed for Dil-LDL uptake, with or without the presence of GPR182 mAb. (F) HAECs were assessed for Dil-LDL transcytosis, with or without the presence of GPR182 mAb. Dil-labeled LDL was added into the inserts at 37°C. After overnight incubation, supernatant from the lower chambers was collected, and fluorescence was quantified using a fluorescence plate reader.

We next assessed GPR182-mediated LDL uptake in human aortic ECs (HAECs). Cultured HAECs constitutively expressed GPR182, which was predominantly localized intracellularly (**Figure S4C**). GPR182 inhibition with a GPR182 blocking mAb (clone 1A5) or siRNA silencing significantly reduced Dil-LDL uptake by HAECs (**Figure 4E, S4D**), which was comparable to that observed upon PCSK9 treatment (**Figure 4E**) or silencing of ALK1 or SR-BI (**Figure S4E, S4F**). In PCSK9-treated HAECs, in which surface LDLR is degraded (**Figure S4E**), the presence of GPR182 mAb further reduced LDL endocytosis (**Figure 4E**), indicating that GPR182-mediated LDL uptake is independent of LDLR. In contrast, GPR182 blockade had minimal impact on LDL uptake by HAECs when either SR-BI or ALK1 was silenced (**Figure S4F**), suggesting that GPR182 may share a pathway with ALK1 or SR-BI for LDL endocytosis.

We next examined the involvement of GPR182 in LDL transcytosis using a transwell assay with HAECs. GPR182 blockade significantly inhibited Dil-LDL transcytosis (**Figure 4F**). Consistent with previous reports^6,8^, silencing of ALK1 or SR-BI significantly reduced Dil-LDL transcytosis. Notably, GPR182 blockade did not further reduce LDL transcytosis in HAECs when either ALK1 or SR-BI was silenced (**Figure S4G**), further supporting the possibility that these receptors participate in a common LDL transport pathway in HAECs.

### GPR182 blockade attenuates dietary-induced atherosclerosis

Having established that genetic GPR182 deficiency attenuates atherosclerosis, we next investigated whether blockade of GPR182 could similarly reduce atherosclerotic lesion development. We used the same GPR182-blocking mAb (clone 1A5) that we previously used to inhibit dietary lipid absorption and treat obesity^17^. Administration of GPR182 mAb right before intravenous injection of Dil-labeled LDL markedly reduced LDL accumulation in the aortic wall (**Figure 5A**), directly demonstrating that the antibody blocks arterial LDL uptake *in vivo* and supporting a vessel wall–directed mechanism of action. To test therapeutic efficacy, we treated *Apoe*−/− mice with GPR182 mAb during 10 weeks of HCD feeding. Compared with control-treated mice, GPR182 mAb-treated *Apoe*−/− mice exhibited significantly reduced atherosclerotic lesion burden, as evidenced by smaller aortic lesion area and reduced lipid deposition (**Figure 5B**, **5C**). Consistent with the en face analysis, cross-sectional assessment of the aortic root demonstrated reduced atherosclerotic lesion area in GPR182 mAb-treated mice (**Figure 5D**).

**Figure 5.**
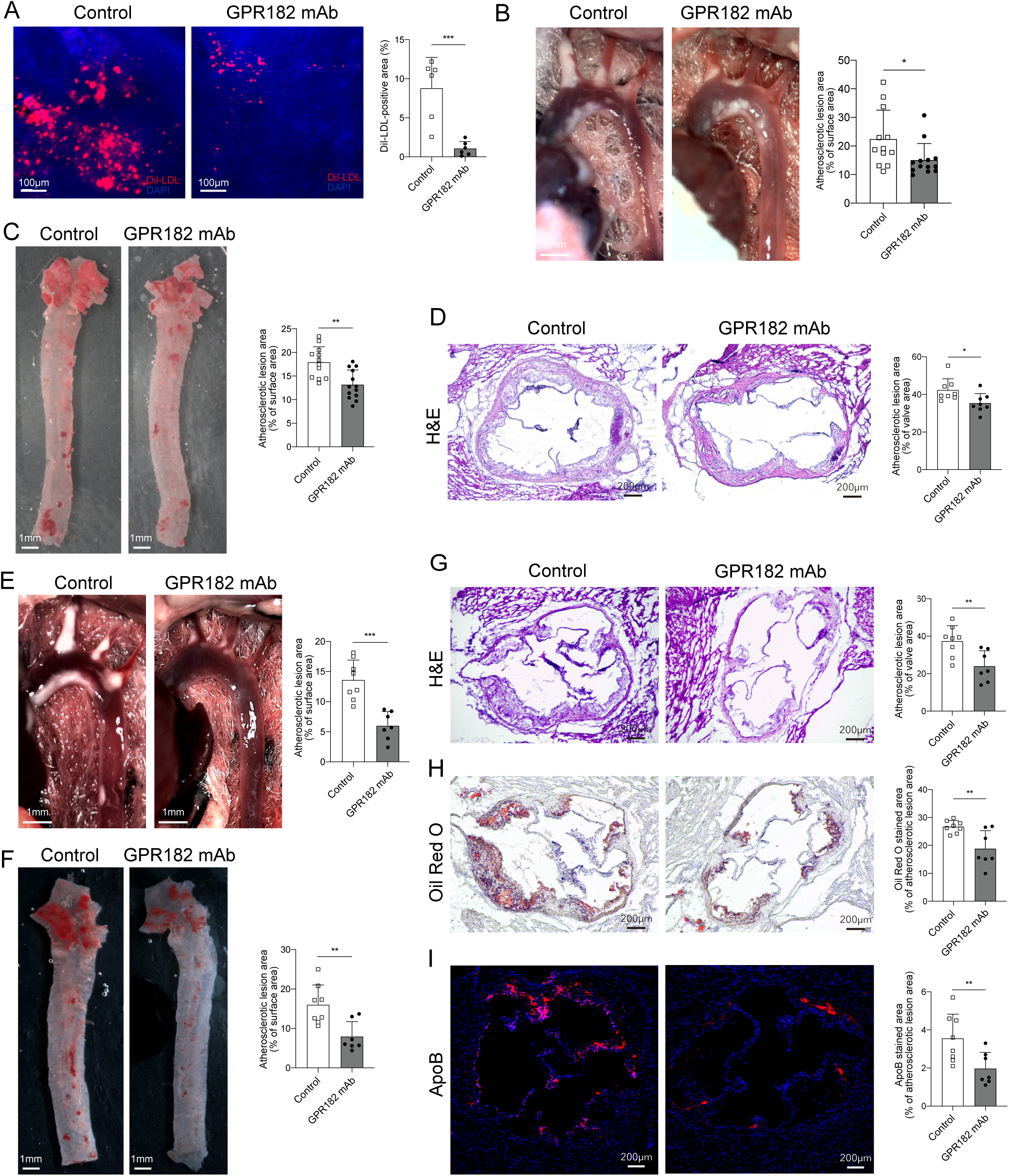
Anti-GPR182 treatment inhibits atherosclerosis progression. (A) Dil-LDL uptake in aorta from WT mice treated with control or GPR182 mAb following intravenous Dil-LDL injection, with quantification of Dil-LDL signal. (B-D) *Apoe*−/− mice under 10 weeks of HCD were treated with control or anti-GPR182 mAb from the start of HCD feeding. (B) Representative in situ aortic arch image of atherosclerotic plaque, with quantification of lesion area. (C) Representative Oil Red O stained en face images of aortas, with quantification of lesion area. (D) H&E-stained sections of aortic root, with lesion area quantification. (E-I) *Ldlr*−/− mice under 10 weeks of HCD were treated control or anti-GPR182 mAb from the beginning of HCD feeding. (E) Representative in situ aortic arch image of atherosclerotic plaque, with quantification of lesion area. (F) Representative Oil Red O stained en face images of aortas, with quantification of lesion area. (G) H&E-stained sections of aortic root, with lesion area quantification. (H) Oil Red O staining of aortic root, with lipid content quantification. (I) ApoB staining in aortic root lesions, with quantification of ApoB-positive area.

We then assessed the atheroprotective effect of GPR182 mAb in *Ldlr*−/− mice under HCD, another widely used mouse model of hypercholesterolemia-driven atherosclerosis^22^. *Ldlr*−/− mice fed an HCD for 10 weeks developed substantial aortic atherosclerosis, though lesion burden was less pronounced than that observed in *Apoe*−/− mice. GPR182 mAb treatment markedly reduced aortic lesion formation and lipid accumulation in *Ldlr*−/− mice (**Figure 5E**, **5F**). Cross-sectional analysis of the aortic root further demonstrated reduced lesion area in GPR182 mAb-treated mice (**Figure 5G**). Plaque composition analysis showed significantly reduced lipid accumulation in lesions from GPR182 mAb-treated mice (**Figure 5H**, **5I**). Like *Gpr182*−/− mice, the treatment of GPR182 mAb had no impact on circulating cholesterol levels in both disease models (**Table S2**). These findings indicate that GPR182 blockade attenuates atherosclerosis independent of any change in circulating cholesterol, including the selective HDL-C increase previously reported in germline Gpr182−/− mice. Together, our studies support GPR182 blockade as a viable approach for treating atherosclerosis.

## Discussion

Our previous publication has established GPR182 as a receptor mediating dietary lipid absorption in the small intestine^17^. In this study, we found that GPR182 is expressed in the aortic endothelium and mediates LDL deposition during the development of atherosclerosis. Genetic GPR182 deletion or blockade with a mAb inhibits atherosclerosis progression without affecting systemic cholesterol. Our study thus reveals a role for GPR182 in endogenous lipoprotein transport and supports receptor-mediated transcytosis as a mechanism for LDL deposition during atherosclerosis.

It has only recently been recognized that LDL entry into the endothelium during the development of atherosclerosis involves receptor-mediated transcytosis ^23^ ^24^. SR-BI and ALK1 are two major surface receptors that have been documented to facilitate LDL deposition in the arterial wall ^9^. Our study here has identified GPR182 as another receptor mediating LDL uptake in the aorta. The *in vitro* studies in HAECs suggest that these three receptors may mediate LDL uptake via a shared pathway. LDL transcytosis in ECs involves caveolin-1-dependent caveolar transcytosis^25^ ^26^, therefore, whether GPR182 mediates LDL endocytosis and transcytosis through the same transport route remains to be determined. GPR182 also functions as an ACKR that scavenges chemokines for degradation through receptor-mediated internalization^11,12^. It thus remains unclear how GPR182 engages distinct ligands and directs lipoproteins and chemokines to different trafficking pathways remains unclear. Defining these mechanisms may reveal how GPR182 coordinates lipoprotein transport and chemokine scavenging in the vascular endothelium.

Our study supports GPR182 as a more feasible therapeutic target in LDL transcytosis in atherosclerosis compared with SR-BI and ALK1. Although SR-BI contributes to endothelial LDL transcytosis, it is broadly expressed, particularly in the liver, where it has a critical role in reverse cholesterol transport^27^. Accordingly, whole-body deletion of SR-BI in mice markedly elevates circulating cholesterol and increases susceptibility to atherosclerosis^28^. Consistent with these observations, human genetic studies have linked *SCARB1* variants to altered HDL-cholesterol levels and increased risk of coronary artery disease^29^ ^30^. Thus, systemic inhibition of SR-BI is not viable and targeting its activity specifically in AECs would be necessary. Targeting ALK1 brings a similarly difficult challenge, as perturbation of ALK1-dependent bone morphogenetic protein (BMP) signaling could have broad physiological consequences, particularly given its essential role in vascular development and physiological angiogenesis^31^ ^32^. In contrast, whole-body GPR182-knockout mice are viable and healthy. Systemic GPR182 blockade with our mAb reduces local LDL deposition in the aortic wall and attenuates atherosclerosis despite having minimal impact on circulating cholesterol levels. Because GPR182 recognizes multiple classes of lipoproteins, its blockade may also limit arterial deposition of TAG-rich lipoprotein (TRL) remnants, which have been implicated in promoting atherosclerosis^33^. Furthermore, GPR182 ablation in our atherosclerosis models does not increase proinflammatory chemokine levels or immune-cell infiltration in atherosclerotic plaques, suggesting that the contribution of GPR182 to arterial lipoprotein deposition predominates over its function as a chemokine scavenger. Nevertheless, selectively disrupting the lipoprotein-uptake function of GPR182 while preserving its chemokine-scavenging activity may further improve the anti-atherosclerotic efficacy.

Although *Gpr182*−/− mice exhibit impaired intestinal lipid absorption, their circulating cholesterol levels are slightly elevated on a regular chow diet ^17^, reflecting a selective increase in HDL cholesterol, which implicates GPR182 in endogenous lipoprotein metabolism. Our finding that GPR182 promotes lipid deposition in the aortic wall supports this. Interestingly, GPR182 deficiency does not alter circulating cholesterol levels in *Apoe*−/− mice or AAV8-PCSK9-transduced mice fed an HCD. Similarly, GPR182 blockade with a mAb has no impact on circulating cholesterol levels in either *Apoe*−/− or *Ldlr*−/− mice under HCD. Collectively, these findings suggest that the effects of GPR182 on exogenous and endogenous lipoprotein transport may counterbalance each other, resulting in minimal net changes in systemic cholesterol levels under hypercholesterolemic conditions; this model remains to be tested directly, for example, with tissue-specific deletion. Besides LECs, GPR182 is broadly expressed in ECs across multiple organs, including liver sinusoidal endothelial cells (LSECs) ^14,34^. Loss of GPR182 in these ECs, particularly LSECs, could impair the clearance of circulating cholesterol-containing lipoproteins, thereby offsetting the reduction of intestinal lipid absorption caused by GPR182 deficiency.

In conclusion, our study reveals a local effect of GPR182 in mediating lipid deposition in the arteries. Targeting GPR182 offers a new therapeutic target for atherosclerosis without affecting circulating cholesterol.

## Methods

### Mice

All mice in C57BL/6J background were breed and housed at the animal facility of the University of Colorado Anschutz Medical Campus under pathogen-free conditions. All animal care procedures and experiments have been approved by the Institutional Animal care and Use Committee at the University of Colorado Anschutz Medical Campus (AMC) (Aurora, CO). Only male mice were used for AAV8-mPCSK9-induced atherosclerosis. For other experiments, both male and female mice were used. C57BL/6, *Apoe*−/−, *Ldlr*−/− mice were originally purchased from the Jackson laboratory. *Gpr182*−/− mice and LEC-specific GPR182 knockout mice (*Gpr182*^ΔLEC^) were generated and described previously^17^. To elucidate the role of GPR182 in atherosclerosis, we generated *Gpr182*−/−*Apoe*−/− mice by crossing *Gpr182*−/− mice with *Apoe*−/− mice. AEC-specific GPR182 knockout mice (*Gpr182*^ΔAEC^) were generated by crossing *Gpr182^fl/fl^* mice with Bmx-CreERT2 mice (rederived by the Genetically Engineered Murine Model Core, University of Colorado AMC, Aurora, CO).

### Reagents and antibodies

Serum cholesterol and TAG concentrations were measured following the manufacturer’s instructions of colorimetric assay kits (Abcam, #ab65390, #ab65359, #ab65336). The following siRNAs and sgRNA were used: GPR182 siRNA, ALK1 siRNA (Dharmacon, D-005303-06), and SR-BI-targeting sgRNA (Synthego). The following antibodies were used for immunofluorescence staining: human CD31 (Cell Signaling, #3528T), human GPR182 (Sigma, #HPA027037), human ALK1 (R&D Systems, #AF-370-SP), mouse GPR182 (recombinant 1A5 chimeric antibody, mVHL-hIgG1κ), mouse ApoB (Proteintech, #20578-1-AP), mouse CD31 (BioLegend, #102408), mouse CD45.2 (BioLegend, #109808), mouse CD4 (BioLegend, #100516), mouse F4/80 (Cell Signaling, #30325). The following antibodies were used for flow cytometry: mouse CD3 (BioLegend, #100312), mouse CD4 (BioLegend, #100449), mouse CD11b (BioLegend, #101208), mouse CD45 (BioLegend, #103122), mouse F4/80 (BioLegend, #123147), mouse CD31 (BioLegend, #102408), mouse GPR182 (clone 1A5), human ALK1 (R&D Systems, #AF-370-SP), human SR-BI (BioLegend, #363203), human LDLR (BD Biosciences, #565653), and viability dye Ghost Dye™ Red 780 (Tonbo Biosciences, #13-0865-T100).

### Mouse models of atherosclerosis

*Apoe*−/− mice and *Gpr182*−/−*Apoe*−/− mice were maintained on regular chow diet until they reached 10 months or older. For food induced atherosclerosis, *Apoe*−/− mice and *Gpr182*−/−*Apoe*−/− mice at 9-11weeks old were fed a high-cholesterol (1.25%) atherogenic diet (HCD) (#D12108C; Research Diets, New Brunswick, NJ, USA) for 10 weeks. For therapeutic assessment, *Apoe*−/− and *Ldlr*−/− mice were fed with HCD for 10 weeks. GPR182 mAb treatment was initiated simultaneously with the onset of HCD. Each mouse receives 200 μg GPR182 mAb intraperitoneally three times a week till analysis. For AAV8-mPCSK9-induced atherosclerosis, 1×10¹¹ vector genome copies of AAV8/D377Y-mPCSK9 (Addgene plasmid: #58376) were intravenously injected to young male mice to increase plasma LDL levels. WT B6, *Gpr182*−/−, *Gpr182*^fl/fl^, *Gpr182*^ΔLEC^, and *Gpr182*^ΔAEC^ mice were used for the experiments. Immediately after AAV injection, mice were switched from a chow diet to HCD. Two weeks after injection, total cholesterol levels in the serum were measured to confirm successful induction of hypercholesterolemia, and only mice with elevated cholesterol levels were maintained on HCD for 12 weeks. Mouse body weights were monitored weekly. After HCD feeding, mice were euthanized and macroscopic evaluation of atherosclerotic lesions was performed from the aortic root, especially the aortic arch. The heart, aorta from the aortic root through the thoracic region, and serum were collected for further analysis.

### Quantification of atherosclerotic lesions

Atherosclerotic lesion area was assessed by imaging of the aorta from the aortic root through the thoracic region, and oil red O staining of en face aorta (Sigma-Aldrich, St. Louis, MO, USA). For en face aorta staining, isolated aorta was carefully removed off the fat and connective tissues before fixed with 10% formalin over night at room temperature. After fixing, the aorta was opened longitudinally from the aortic root to the thoracic aorta. Oil red O was used for atherosclerotic plaque staining. After the staining, aortic images were captured with a digital camera and the total surface and entire lesion areas were measured by Image J (National Institutes of Health, Bethesda, MD, USA).

### Immunohistochemistry and immunofluorescent staining

Samples of hearts including aortic root were embedded in optimum cutting temperature (OCT) compound (Leica Microsystems GmbH, Wetzler, Germany) and 8 µm thick frozen sections were prepared. Formalin-Fixed Paraffin-Embedded (FFPE) tissue blocks and sections were processed and prepared by the Pathology Shared Resource at the University of Colorado AMC. FFPE sections were rehydrated and treated in Tris-EDTA buffer (pH 9.0 or pH 5.0) at 110°C for 15 minutes before proceeding with staining steps. For immunohistochemistry staining, the sections were stained with hematoxylin and eosin staining (H&E), Picro Sirius Red (Abcam, Cambridge MA, USA), Oil Red O staining as previously described. Aortic root area, lesion area, oil area, and collagen area analysis were conducted using the ImageJ software. For Immunofluorescent staining, tissues were blocked with 2.5% goat serum and incubated with primary antibodies at the appropriate concentration overnight at 4°C. Secondary antibodies were incubated for 1 hour, followed by counterstaining for 7 minutes at room temperature. After clearing and mounting with Fluoromount-G™ Mounting Medium (Thermo Fisher), immunofluorescent images were captured with ZEISS Axio Observer and analyzed with SlideBook software (Version 6, Intelligent Imaging Inc).

### Flow cytometry

Single-cell suspensions were prepared by cutting aortic tissues into small pieces and were digested with 50 μg/mL Liberase (Roche Diagnostics Corporation, Indianapolis, IN, USA) in PBS at 37°C for 1hour. After digestion, single-cell suspensions were collected by passing them through a 70 μm cell strainer (Fisherbrand, Cat# 22363548). Cells were then washed with culture medium and were incubated with LEAF anti-mouse CD16/32 (anti-FcγRIII/II receptor, clone 93) for blocking prior to staining. Surface markers were stained with fluorophore-conjugated antibodies for 30 minutes at 4°C. Viability staining was performed with Ghost Dye Red 780 (Tonbo Biosciences, #13-0865-T100). Flow cytometric analysis was conducted using a Northern Lights or Aurora cell analyzer (Cytek Biosciences), and data were analyzed with FlowJo software (version 10.9.0, Tree Star).

### FPLC analysis

For FPLC analysis, serum samples (400 μL) were fractionated by size-exclusion chromatography using two Superose 6 columns connected in series, as previously described^17^. During elution, absorbance was monitored at 280 nm. Fractions corresponding to VLDL, LDL, and HDL were collected, and cholesterol concentrations in each fraction were measured using a cholesterol assay kit (Abcam, #ab65390) according to the manufacturer’s instructions.

### LDL endocytosis in *vivo*

WT and *Gpr182*−/− mice were injected via tail vein with 100 μl of Dil-LDL (300 μg/mouse). Thirty minutes later, mice were euthanized and aortae were perfused with 10 ml of PBS, followed by 10 ml of 4% paraformaldehyde fixation. Mouse aorta (atheroprone area; low curvature of aortic arch around branch) were dissected and mounted. Images were taken by ZEISS Axio Observer. For immunofluorescent analysis, aortic tissues from WT mice were embedded in OCT compound and cryosectioned. Sections were subjected to immunofluorescent staining using antibodies against GPR182 and CD31 to assess their colocalization with Dil-LDL signal.

### Endothelial permeability assay *in vivo*

Evans blue (1.0%, 100 μL; Sigma-Aldrich, E2129) was injected via the tail vein. After 30 minutes, mice were euthanized, and the heart and aorta were carefully dissected under a microscope. Aortas were dried and weighed, and Evans blue was extracted by incubation in formamide at 60°C for 24 hours. Absorbance was measured at 620 nm. A standard curve generated using Evans blue was used to calculate the total amount of extracted dye, which was adjusted to the weight of isolated aortas.

### LDL endocytosis *in vitro*

Human aortic endothelial cells (HAECs), purchased from Lonza (#CC-2535), were cultured in Endothelial Cell Growth Basal Medium-2 with Endothelial SingleQuots^®^ Kit (Lonza, #CC-3156, #CC-4176). For LDL endocytosis assay, HAECs were fasted for 1 hour before incubated with Dil-LDL (5μg/ml; Kalen Biomedical, #770230-9) at 37°C for 30 minutes. In some assays, HAECs were incubated with PCSK9 (10μg/ml) at 37°C overnight to reduce LDLR expression, which was confirmed by flow cytometry. To assess the involvement of GPR182, cells were treated with GPR182 mAb (clone 1A5) or transfected with GPR182 siRNA. For receptor interaction studies, HAECs were transfected with ALK1 siRNA or SR-BI-targeting sgRNA. Silencing of ALK1 and SR-BI in HAECs was confirmed by flow cytometry. Cells were subsequently treated with GPR182 mAb (clone 1A5) where indicated, followed by incubation with Dil-LDL for 30 minutes. Cells were stained with WGA and spectral DAPI before analysis. Images were taken by ZEISS Axio Observer.

### LDL transcytosis *in vitro*

LDL transcytosis was evaluated using a modified transwell assay based on a previously reported method^17^. HAECs were seeded onto collagen I (BD-Bioscience) coated 0.4-μm pore PET Transwell inserts (Corning, #3610). LDL transcytosis was assessed 48 hours after seeding, when a confluent endothelial monolayer had formed. For receptor interaction studies, HAECs were transfected with ALK1 siRNA or SR-BI-targeting sgRNA before the transcytosis assay. Fluorescein isothiocyanate (FITC)-dextran (30 μg/mL, molecular weight 3,000; Invitrogen) was used to assess paracellular transport. Cells were incubated simultaneously for overnight with FITC–dextran and 50 μg/ml Dil–LDL added to the upper chamber. In experiments using GPR182 blockade, cells were pretreated with GPR182 mAb (clone 1A5, 50 μg/mL) in the upper chamber for 1 hour before the addition of Dil-LDL and FITC-dextran. After overnight incubation at 37°C, fluorescence intensities of Dil and FITC in the lower chamber were measured using a fluorometer (Infinite M Plex, Tecan). FITC–dextran transport was approximately 3 %, confirming minimal paracellular leakage for the endothelial monolayer.

### Chemokine quantification

Aortic samples were homogenized in 1 mL of 1× PBS containing protease inhibitors using a Fisherbrand™ 150 handheld homogenizer. The homogenates were centrifuged at 14,000 × g for 15 minutes to collect the supernatants. Chemokines in the supernatants were subsequently quantified using the LEGENDplex Mouse Proinflammatory Chemokine Panel (BioLegend) and analyzed according to the manufacturer’s instructions using a Beckman Coulter CytoFlex S. Total protein concentration in each sample was measured using the Bradford assay with Coomassie Blue (Bio-Rad) and used to normalize chemokine levels to total protein content in the supernatants.

### Gene expression analysis in human atherosclerosis

To evaluate GPR182 expression in human atherosclerosis, the publicly available gene expression dataset GSE43292 was obtained from GEO (https://www.ncbi.nlm.nih.gov/geo/). The dataset contains paired atherosclerotic and non-atherosclerotic aortic tissues.

### Single-cell RNA-sequencing analysis

Publicly available single-cell RNA-sequencing data from the Tabula Muris Senis dataset were downloaded from the Tabula Muris Senis data portal (https://tabula-muris-senis.ds.czbiohub.org/) and analyzed to assess *Gpr182* expression across annotated cell populations in mouse aorta and heart. The data were analyzed and visualized using Scanpy (version 1.9.3).

### Statistical analysis

All the experiments were repeated independently. The data were presented as mean ± standard deviation (SD). Unpaired two-tailed Student’s t test was used to compare the means of two groups. Two-way ANOVA with Bonferroni’s correction for multiple comparison was used to compare groups over time by repeated measures. All P values less than 0.05 were deemed to be significant. GraphPad Prism 9.0 (GraphPad Software, Inc., La Jolla, CA, USA) was used for all statistical analysis and to generate figures.

## Author contributions

TT and YZ designed the study; TT and SD conducted experiments and acquired the data for most of the study; ZS, YS, JH, DPF, YG, YH, assisted with experiments and data acquirement; YZ, RDS and MS provided with resource and assistance; YZ conceived and supervised the project; YZ, TT, and SD wrote the manuscript; all authors read and approved the final manuscript.

## Acknowledgments

We thank the Functional Genomics Shared Resource and the Pathology Shared Resource, supported by the Cancer Center Support Grant at the University of Colorado (P30CA046934), for resources and services used in this study. We thank the Mouse Genetics Core Facility at the University of Colorado Anschutz Medical Campus for sperm rederivation.

