## Supplemental Figures for "GPR182 promotes atherosclerosis via facilitating arterial lipid deposition"

**Table S1 Serum TC and TAG levels in *Apoe*$^{-/-}$ and *Gpr182*$^{-/-}$*Apoe*$^{-/-}$ mice**

| | | *Apoe*$^{-/-}$ | *Gpr182*$^{-/-}$*Apoe*$^{-/-}$ | p value |
|---|---|---|---|---|
| **TC (mg/dL)** | Pre-HCD male | 801.3 ± 156.0 (8) | 930.7 ± 184.4 (7) | 0.1642 |
| | Pre-HCD female | 509.4 ± 136.9 (8) | 535.0 ± 144.2 (8) | 0.7209 |
| | Post-HCD male | 2947.5 ± 749.4 (8) | 2731.4 ± 625.2 (7) | 0.5583 |
| | Post-HCD female | 2548.6 ± 460.6 (7) | 2406.7 ± 264.3 (6) | 0.5202 |
| | 10-month-old male | 1496.7 ± 65.1 (3) | 1570.0 ± 195.2 (3) | 0.5704 |
| | 10-month-old female | 785.0 ± 222.2 (3) | 911.7 ± 118.6 (3) | 0.4329 |
| **TAG (mg/dL)** | Pre-HCD male | 102.4 ± 34.4 (8) | 106.3 ± 20.6 (7) | 0.7975 |
| | Pre-HCD female | 67.0 ± 8.0 (8) | 64.4 ± 17.1 (8) | 0.6994 |
| | Post-HCD male | 92.1 ± 29.2 (8) | 95.7 ± 32.2 (7) | 0.8245 |
| | Post-HCD female | 62.0 ± 16.2 (7) | 64.5 ± 21.2 (6) | 0.8140 |
| | 10-month-old male | 84.3 ± 26.5 (3) | 85.0 ± 14.0 (3) | 0.9711 |
| | 10-month-old female | 39.3 ± 5.0 (3) | 40.7 ± 2.3 (3) | 0.6981 |

*TC* = Total cholesterol; *TAG* = Triacylglycerol;

**Table S2 Serum TC levels in *Apoe*$^{-/-}$ and *Ldlr*$^{-/-}$ mice treated with GPR182 mAb**

| | | Control | GPR182 mAb | p value |
|---|---|---|---|---|
| **TC (mg/dL)** | *Apoe*$^{-/-}$ Post-HCD male | 1806.7 ± 402.7 (6) | 1793.3 ± 285.6 (6) | 0.9486 |
| | *Ldlr*$^{-/-}$ Post-HCD male | 2300.0 ± 431.4 (6) | 2093.3 ± 483.1 (6) | 0.4525 |

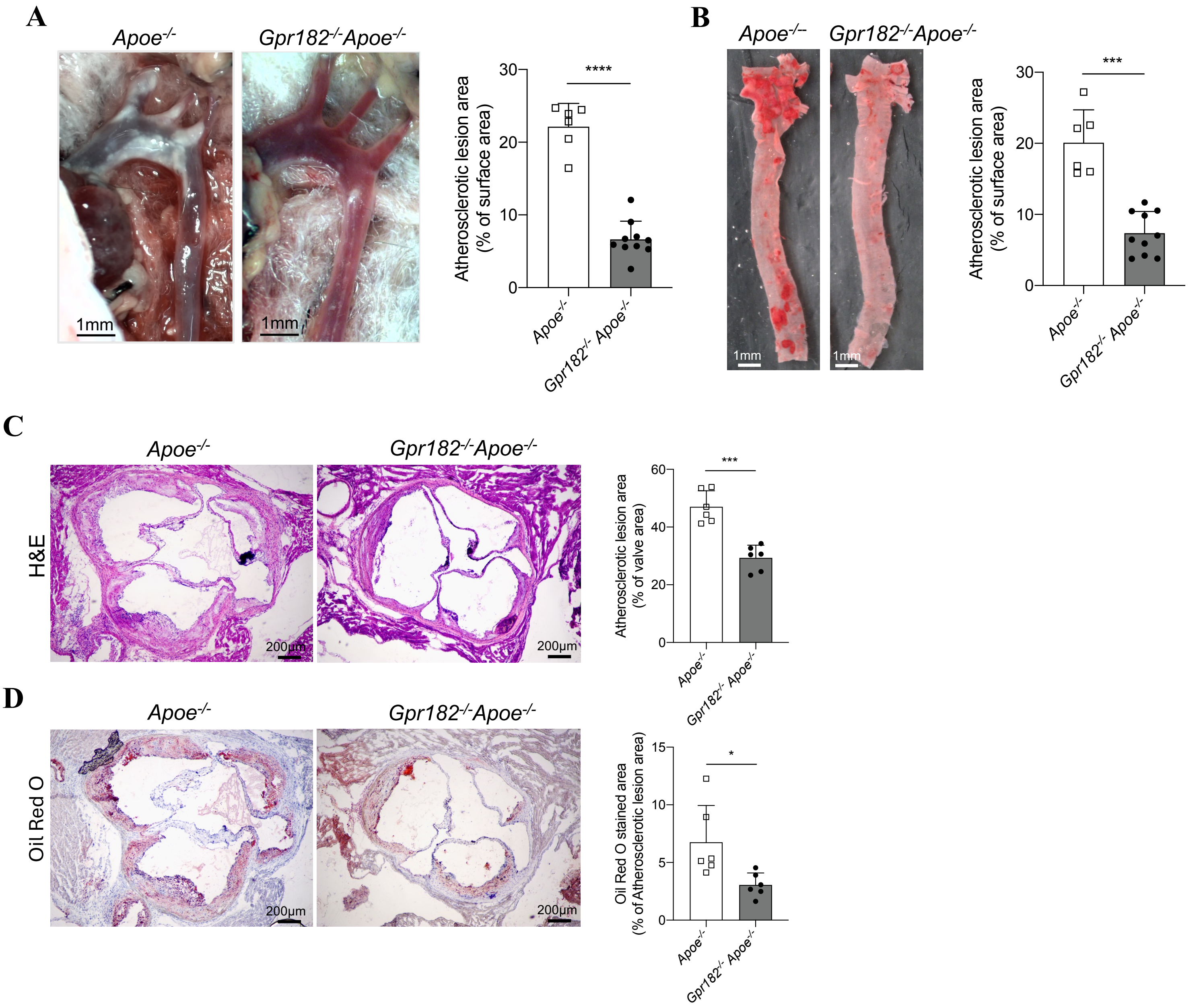


**Supplementary Figure 1 GPR182 ablation attenuates age-associated atherosclerosis in *Apoe*$^{-/-}$ mice.** (A) In situ images of atherosclerotic plaques in the aortic arch, with quantification of lesion area. (B) En face Oil Red O staining of thoracic aortas, with quantification of lesion area. (C) H&E-stained sections of aortic root, with lesion area quantification. (D) Oil Red O staining of aortic root, with lipid content quantification.

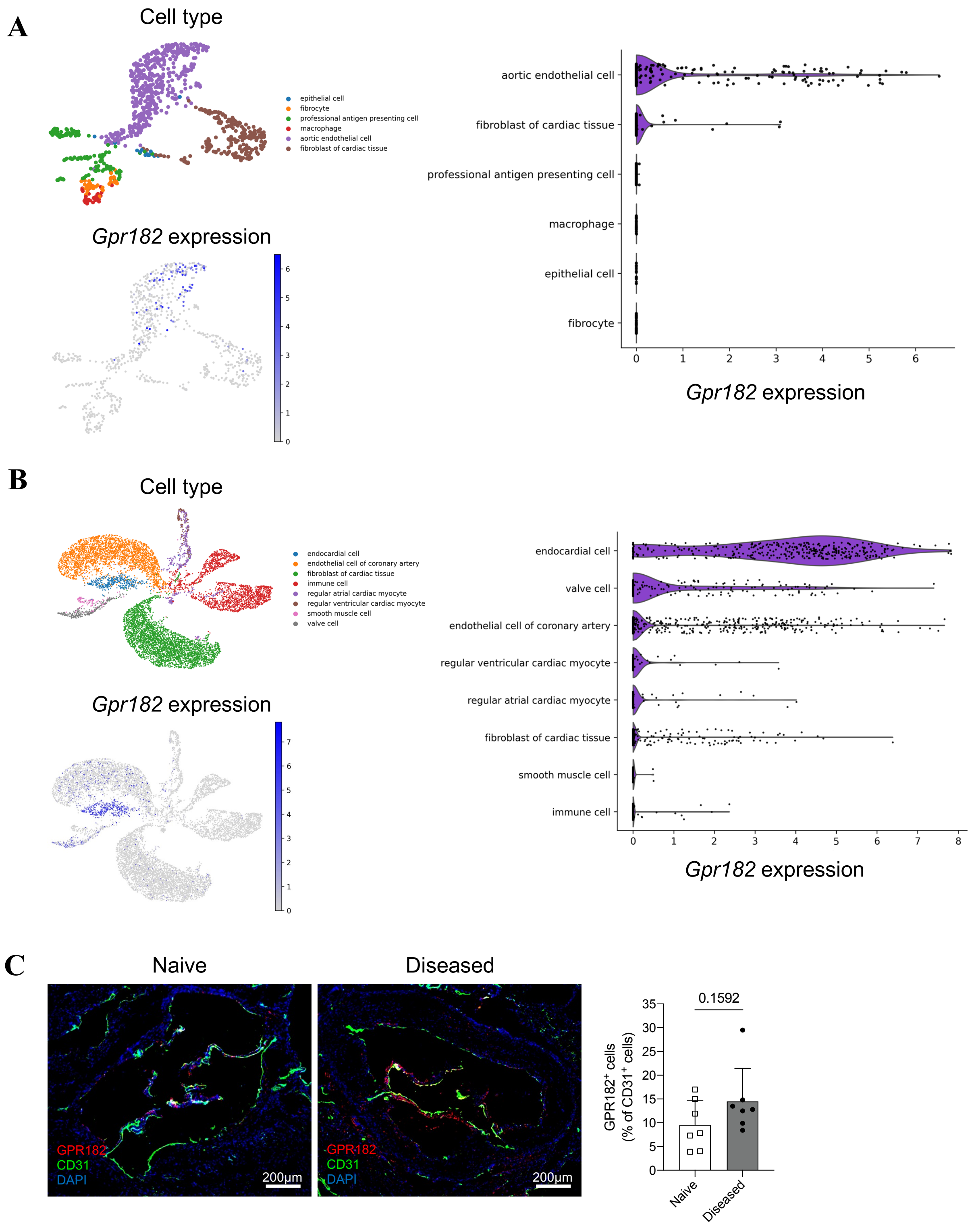


**Supplementary Figure 2 GPR182 is abundantly expressed in cardiac ECs.** (A, B) The Tabula Muris single-cell RNA-sequencing dataset was reanalyzed for *Gpr182* expression in mouse aorta (A) and heart (B). (C) GPR182 expression in aortic tissues of naïve WT B6 and *Apoe*-/- mice with atherosclerosis (Diseased) was assessed by immunofluorescence staining.

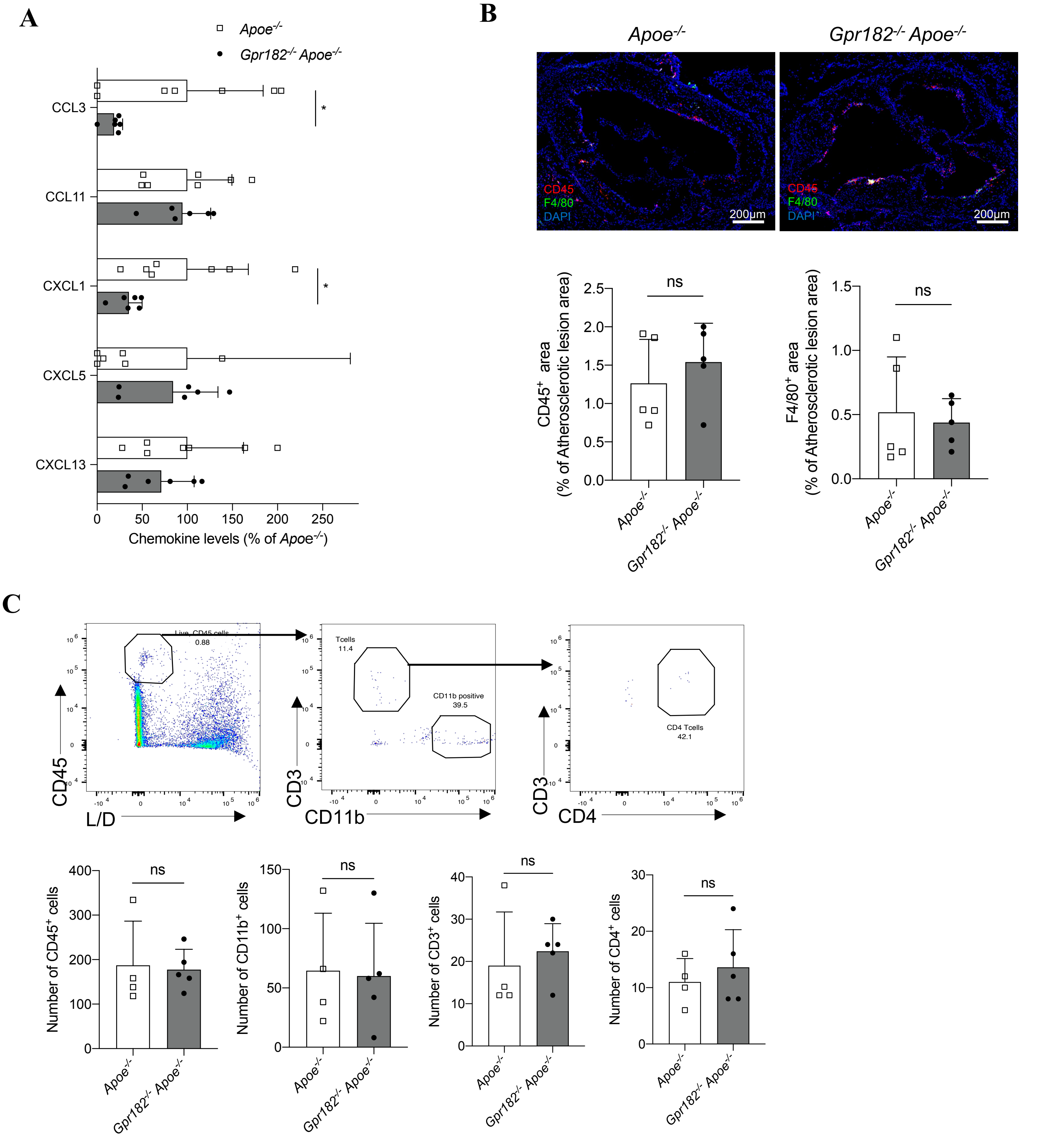


**Supplementary Figure 3 GPR182 ablation does not affect immune cell accumulation in atherosclerotic plagues.** (A) Chemokine levels in aortas from ApoE$^{-/-}$ and GPR182$^{-/-}$ApoE$^{-/-}$ mice. (B) CD45 and F4/80 staining in aorta plaques, with quantification of CD45$^{+}$ and F4/80$^{+}$ cells. (C) Immune cell populations in aorta assessed by flow cytometry, with quantification of CD45$^{+}$, CD11b$^{+}$, CD3$^{+}$, and CD4$^{+}$ cells.

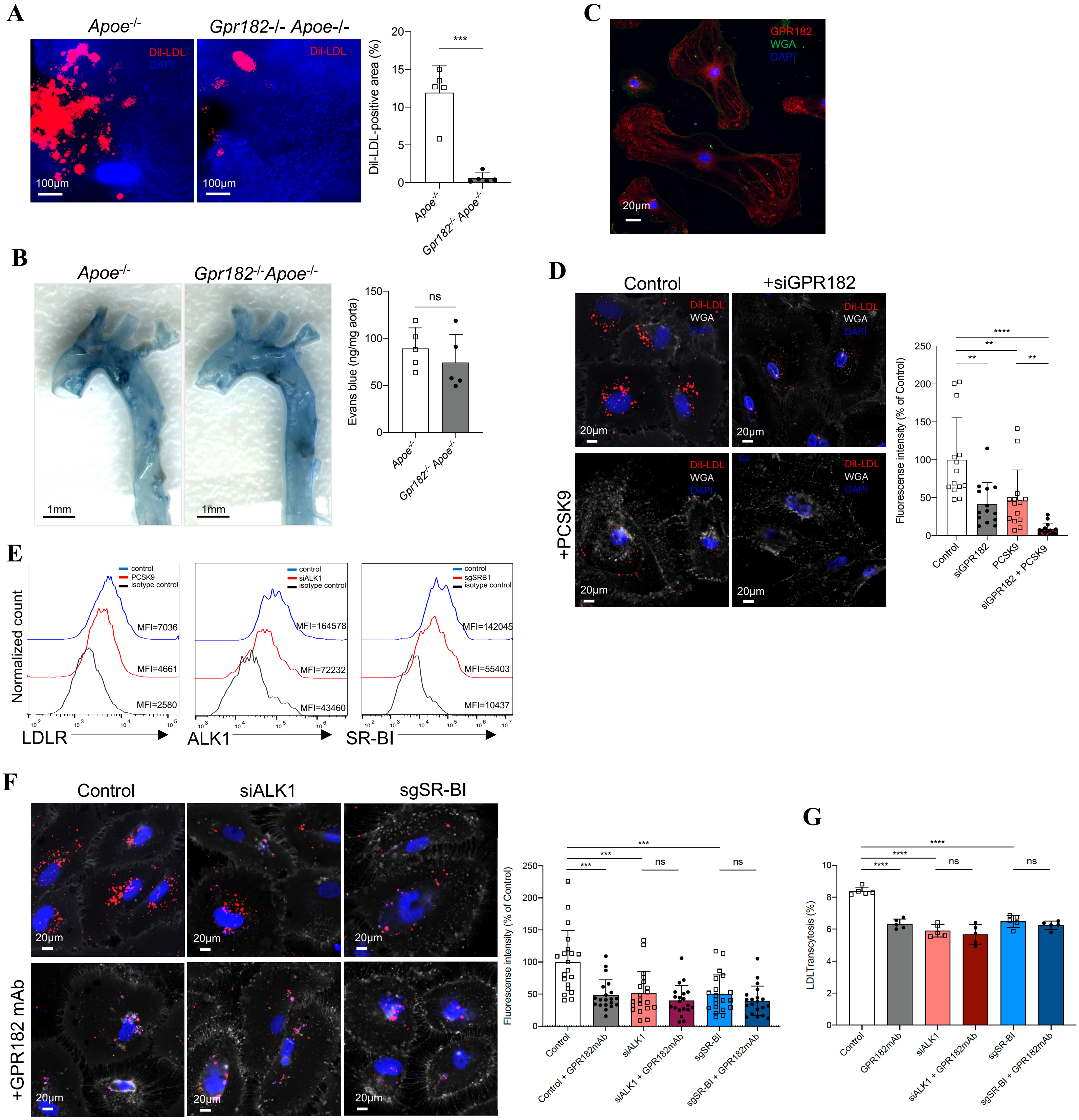


**Supplementary Figure 4 GPR182 mediates LDL endocytosis and transcytosis in arteria ECs.** (A) Dil-LDL deposit in the aorta was determined after intravenously injected into *Apoe-/-* and *Gpr182-/-Apoe-/-* mice. (B) Evens blue signal was examined in the aorta after *Apoe-/-* and *Gpr182-/-Apoe-/-* mice were intravenously injected with Evens blue dye. (C) GPR182 expression in cultured HAECs was detected by IF staining. (D) HAECs and HAECs knocked down of GPR182, with or without PCSK9 treatment, were assessed for DiI-LDL uptake. (E) The reduction in LDLR expression following PCSK9 treatment and the knockdown of ALK1 and SR-BI following gene silencing in HAECs were confirmed by flow cytometry. (F) HAECs with or without ALK1 or SR-BI silencing were assessed for Dil-LDL uptake in the presence or absence of GPR182 mAb, as indicated. Representaptive images were indicated. (G) HAECs with or without ALK1 or SR-BI silencing were assessed for DiI-LDL transcytosis in the presence or absence of GPR182 mAb, as indicated.
